# Membrane-targeting antimicrobials trigger lysis in *Bacillus subtilis* by disturbing the MreB-dependent regulation of peptidoglycan hydrolases

**DOI:** 10.64898/2026.05.01.722135

**Authors:** Kenneth H. Seistrup, Alan Koh, Henrik Strahl

## Abstract

Membrane-targeting antimicrobials are generally assumed to kill bacteria through bacteriolysis induced by the permeabilisation of the cytoplasmic membrane. This model relies on the notion that bacteriolysis is the direct cellular manifestation of the membrane-disruptive (membranolytic) activity of an antibacterial compound. However, it underappreciates the key role of peptidoglycan hydrolases in bacteriolysis. Using the Gram-positive model organism *Bacillus subtilis*, we demonstrate that the bacteriolytic activity of membrane-targeting antimicrobials arises from the misregulation of peptidoglycan hydrolases, regardless of whether they induce large membrane pores or trigger more subtle membrane disturbances such as depolarisation. Contrary to previous models, the autolysis of *B. subtilis* induced by membrane depolarising compounds does not depend on pH changes associated with the cell surface. Instead, the autolytic process is triggered by membrane depolarisation-dependent dissociation of the bacterial actin homolog MreB, a key spatial coordinator of cell wall synthesis in rod-shaped bacteria, from the cytoplasmic membrane. These findings provide valuable insights into the cellular pathways involved in autolysis, highlight the challenges in distinguishing between direct and indirect cellular effects of membrane-targeting antibiotics, and improve our understanding of antibiotic-induced bacteriolysis.

## INTRODUCTION

Most bacteria are surrounded by a cell wall sacculus composed of cross-linked peptidoglycan strands, which allows the cell to maintain turgor and a specific cell morphology. During cell elongation and division, the peptidoglycan sacculus must be locally opened to allow insertion of new glycan strands, followed by the degradation of the outermost peptidoglycan layers to enable the expansion of the cell wall, and the septal peptidoglycan must then be cleaved to allow daughter cell separation in Gram-positive bacteria^1–3^. Peptidoglycan remodelling is important for cell differentiation, assembly of envelope-spanning structures such as flagella, and developmental or fratricidal lysis^2,4,5^. These processes are driven by peptidoglycan (PG) hydrolases with partially overlapping activities, including amidases, endopeptidases, lytic transglycosylases, carboxypeptidases, and glucosaminidases^1–3^. While this enzymatic redundancy provides robustness, it also necessitates multi-layered regulatory systems, as inappropriate or excessive cleavage can lead to unintended catastrophic autolysis^1–3^. Unsurprisingly, PG hydrolases are regulated at the level of enzyme activation, spatial localisation as well as abundance^1,4^. Direct control is exerted by cognate regulators that activate or inhibit hydrolases in response to cell cycle cues or envelope status, by small molecules, or by assembly into multi-enzyme complexes where hydrolases are coupled to PG synthases^1,4^. Spatial regulation is achieved through recruitment by scaffolding proteins, association with surface polymers such as teichoic acids, or substrate modification that restricts access of particular hydrolases to defined cell wall regions^1,2^. Finally, transcriptional and post-transcriptional mechanisms adjust hydrolase abundance to growth conditions and stress, often via two-component systems and envelope stress-responsive sigma factors ^1,2^. Together, these regulatory strategies ensure that PG cleavage is tightly coordinated with PG synthesis in both time and space, allowing controlled cell wall expansion while minimising the risk of lethal stress-induced lysis (autolysis).

The PG hydrolases of the Gram-positive model organism *Bacillus subtilis* exhibit typical functional diversity and regulatory complexity^1,2^. While its genome encodes 42 predicted or confirmed PG hydrolases, the presence of only one of the two elongation-linked D,L-endopeptidases (either LytE and CwlO) is needed to support vegetative growth, with inactivation of both causing a lethal block in cell elongation^6–8^. LytE and CwlO cleave peptide crosslinks within the wall to permit insertion of new material during sidewall synthesis, while other autolysins such as LytC, LytD and LytF contribute phenotypically to cell chain separation^2,3,7,9^. As in other bacteria, multiple enzymes contribute to cell envelope breakdown during antibiotic or stress-induced lysis (autolysis), and inactivation of major hydrolases strongly suppresses the autolytic response^2,10–13^.

An important yet mechanistically poorly understood regulatory mechanism in *B. subtilis* is the coupling of PG hydrolase activity to the cytoskeletal machinery that drives cell morphogenesis^14,15^. During cell elongation, the elongasome, built around the actin homologues MreB, Mbl and MreBH and the PG synthase module RodA–PbpA/PbpH, inserts new PG circumferentially along the lateral wall^16–18^. Genetic and cell biological analyses demonstrated that CwlO activity is controlled by the ABC-transporter-like complex FtsEX, and that Mbl is required for proper FtsEX-CwlO function, whereas MreB and MreBH genetically interact with the LytE system^6,8,14^. Additional layers of regulation are imposed by envelope stress pathways, including the extracytoplasmic function (ECF) sigma factors that modulate the expression of autolysins and their regulators^19,20^. Recent research has also shown that *B. subtilis* can directly modulate PG hydrolase activity. For example, changes in Mg^2+^ availability alter PG hydrolase activity and can rescue the otherwise lethal morphology of *mreB* deletion mutants^15^. A similar role in regulating PG hydrolase activity has been demonstrated for teichoic acids in several Gram-positive species, including *B. subtilis*^21–23^.

*B. subtilis* undergoes extensive autolysis in response to conditions and agents that compromise cell membrane integrity or deplete the cell’s energy sources ^24–27^, which implies severe misregulation of PG hydrolase activity. As the common nominator, these autolysis-inducing stresses lead to the collapse of the proton motive force (PMF)^25–27^, and the associated changes in membrane physicochemical properties have been postulated to trigger activation of PG hydrolases that ultimately leads to cell lysis^26^. However, the molecular mechanisms underpinning this lethal cellular process have remained elusive. Here, we demonstrate that, rather than relying on changes in membrane properties and local pH to regulate PG hydrolase activity, de-energisation-induced autolysis is a more complex cellular process that involves Mre-cytoskeleton-mediated regulation of PG hydrolases and is initiated by the delocalisation of MreB to the cytoplasm upon PMF dissipation. Furthermore, we show that lysis induced by antimicrobials that create large membrane pores arises from the same MreB-dependent misregulation of PG hydrolases, thereby challenging our current understanding of how membrane-targeting antimicrobials exert their bacteriolytic effects.

## RESULTS

### *B. subtilis* cell lysis triggered by PMF collapse is autolysin-dependent

To establish experimental conditions to analyse how the dissipation of proton motive force (PMF) induces cell lysis in *B. subtilis*, and to observe the process at the single-cell level, we followed the response of *B. subtilis* grown in LB medium to the proton uncoupler CCCP^28^, which dissipates both components of the PMF, the transmembrane electric potential (ΔΨ) and the transmembrane pH difference (ΔpH). As shown in Fig. 1a, *B. subtili*s growth was rapidly arrested upon addition of CCCP, followed by a delayed cell lysis indicated by a decline in the culture optical density. Fluorescence microscopy using the membrane permeability reporter Sytox Green^29^, a DNA-intercalating fluorophore that is unable to diffuse across intact membranes, confirmed that the cytoplasmic membranes initially remained intact in the presence of CCCP (Fig. 1b). Upon extended incubation, however, the cell membranes became permeabilised as indicated by the strong Sytox Green fluorescent signal, followed by more extensive leakage of the cellular content and untimately cell disintegration, consistent with cell lysis^30^. These experiments thus confirm that dissipating PMF indeed triggers delayed cell lysis in *B. subtilis* under our experimental conditions.

**Figure 1:**
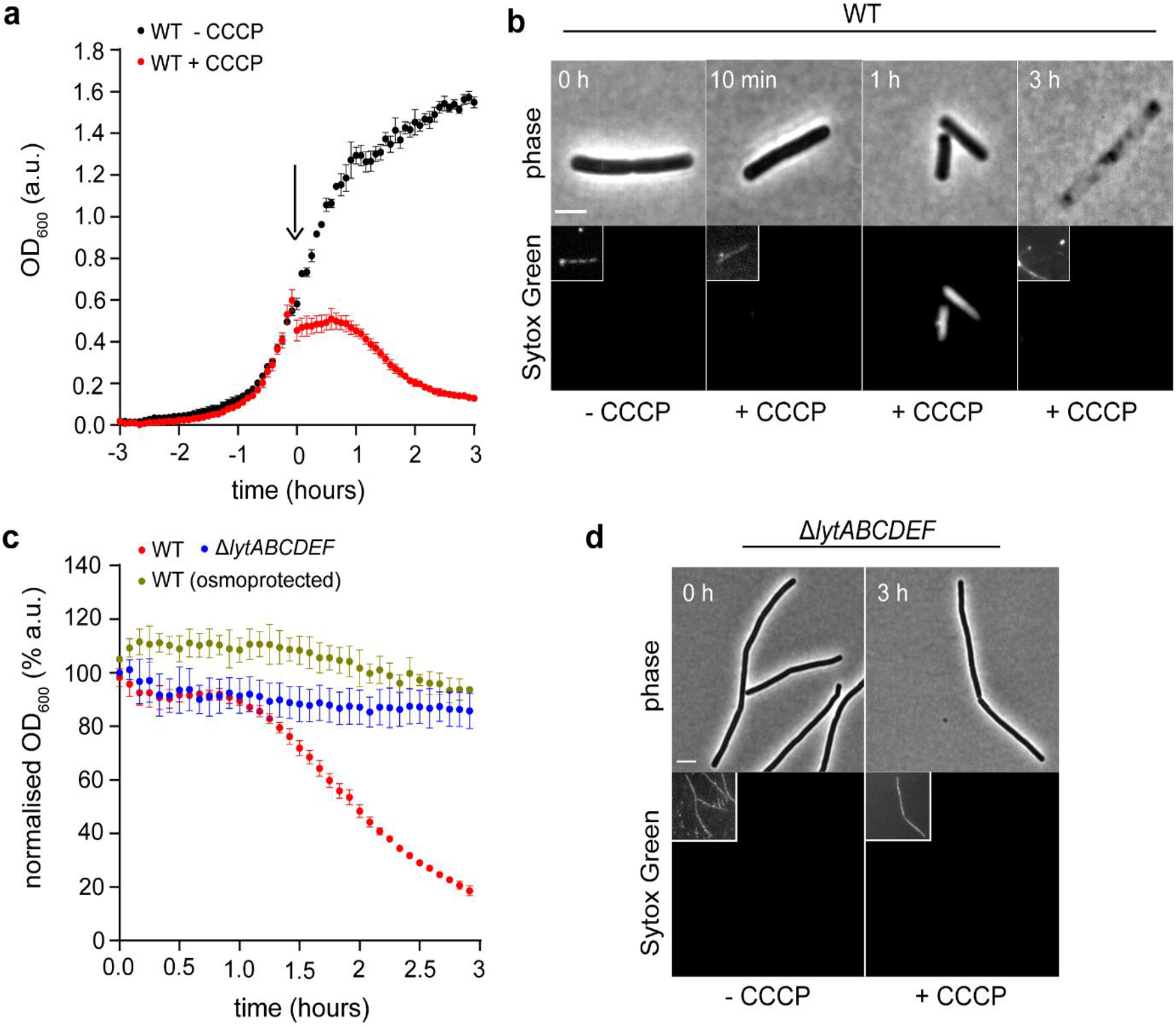
Collapse of PMF triggers autolysis in *Bacillus subtilis*. **(a)** Membrane uncoupler CCCP triggers rapid growth arrest followed by delayed cell lysis in *B. subtilis*. The graph depicts the growth and lysis of *B. subtilis* in LB medium in the absence and addition of CCCP at a time point indicated with an arrow. **(b)** *B. subtilis* cells experience delayed membrane permeabilisation upon incubation with CCCP (1h), followed by cell disintegration at later stages (3h). The phase contrast and fluorescence micrographs depict cells stained with the membrane permeability-indicator Sytox Green, and incubated with CCCP for the times indicated. **(c)** The observed lysis upon incubation with CCCP is autolytic. The graph depicts the kinetics of CCCP-induced cell lysis for wild-type cells in regular LB medium and medium supplemented with 500 mM sucrose, and for cells deficient for several autolysins (Δ*lytABCDEF*). **(d)** CCCP-induced delayed membrane permeabilisation observed for wild-type cells is abolished in cells deficient for several autolysins (Δ*lytABCDEF*). The phase contrast and fluorescence micrographs depict cells stained with the membrane permeability-indicator Sytox Green and incubated with CCCP for 3h. The graphs depict the mean and SD of OD_600_ measurements performed in technical triplicate. The small image inserts serve to visualise cells in image fields that are dark due to very low Sytox Green fluorescence. Scale bar, 3 μm. Strains used: *B. subtilis* 168 (wild-type) and *B. subtilis* KS19 (Δ*lytABCDEF*).

*B. subtilis* genome encodes 42 proteins with either predicted or confirmed peptidoglycan hydrolase activities. However, only one (either LytE or CwlO) is needed for growth and division^28–32^. To confirm that cell lysis observed upon PMF collapse is indeed autolytic, and to analyse the contribution of individual PG hydrolases, we focused on the best characterised peptidoglycan hydrolases LytC, LytD, LytF, LytE and CwlO^31–35^. While strains deficient for individual PG hydrolases exhibited slightly altered lysis kinetics, no individual enzyme was solely responsible for the CCCP-induced cell lysis (Fig. S1a). Similarly, neither the absence of FtsEX complex, which regulates the activity of CwlO^33^, nor lipoteichoic acids, which are involved in autolysis in S*treptococcus pneumoniae*^22^, suppressed the cell lysis (Fig S1a-b). However, deletion of multiple PG hydrolase genes (Δ*lytABCDEF*) strongly suppressed the autolysis induced by CCCP (Fig. 1c-d). This is consistent with previous studies^13,24^ and argues for the involvement of multiple PG hydrolases in the lysis process. However, the Δ*lytABCDEF* strain exhibits a defect in cell separation, resulting in growth as long chains (Fig. S1c). To rule out that this altered cell morphology interferes with our lysis assay, we monitored the CCCP-induced lysis of wild-type cells in a high-osmolarity medium, which should allow the cells to cope with a weakened cell wall without lysis. As with the Δ*lytABCDEF* strain, lysis was suppressed (Fig. 1 c). These findings demonstrate that cell lysis induced by PMF collapse is indeed driven by PG hydrolases and, thus, is autolytic in its nature. Intriguingly, the loss of viability of *B. subtilis* in the presence of CCCP was not suppressed by the lack of autolysis (Fig. S1d). This indicates that lysis is not solely responsible for the bactericidal effect of PMF collapse in *B. subtilis*, and lysis-independent processes such as induced ROS production likely contribute significantly to this effect^36^.

### Induced autolysis is growth phase dependent but not a programmed cellular response

While studying CCCP-induced autolysis, we observed that the process is temperature-dependent, with cells grown at 30°C exhibiting significantly suppressed autolysis (Fig. 2a-b). This is surprising as a change from 37°C to 30°C is unlikely to substantially alter the respiration-dependent changes in cell envelope physicochemical properties that have previously been postulated to activate PG hydrolases upon PMF collapse^26^. In principle, *B. subtilis* cells adapted to faster growth rates at 37°C could exhibit higher PG hydrolase activities, which are required for faster cell wall processing and turnover. To analyse the correlation between growth and lysis, the growth rates were modified by altering the medium carbon sources. While the observed lysis rates varied with media composition, no positive correlation was observed between growth rates and subsequent lysis (Fig. S2). Another growth parameter that should affect cell wall turnover is the growth phase, with non-growing, stationary-phase cells likely exhibiting reduced cell wall turnover. Indeed, CCCP-induced autolysis exhibited a clear growth phase dependency whereby stationary growth phase cells obtained through overnight incubation exhibited no autolytic behaviour, while early logarithmic growth phase cells displayed partial suppression, likely reflecting cells still undergoing phenotypic transition (Fig. 2c).

**Figure 2:**
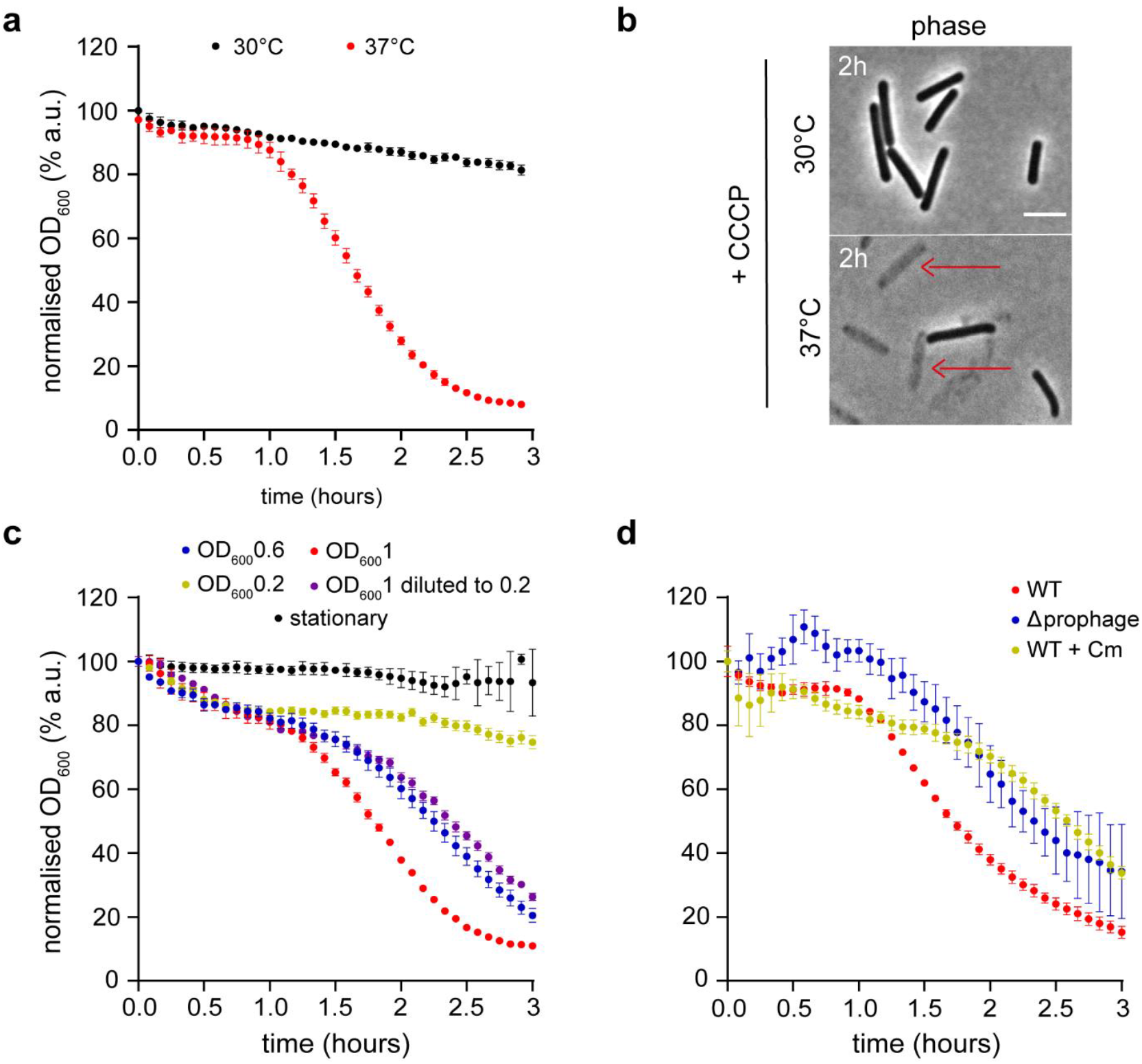
*B. subtilis* autolysis upon PMF collapse is temperature and growth phase-dependent. (**a**) Wild-type cells exhibit a slower rate of CCCP-induced autolysis when grown at 30°C. The graph depicts the kinetics of CCCP-induced autolysis of mid-logarithmic growth phase cells grown and incubated at 30°C and 37°C in LB medium. (**b**) Phase contrast images of wild-type cells showing extensive autolysis upon 2h incubation in the presence of CCCP at 37°C but not at 30°C. Some of the lysed cells are indicated with red arrows. (**c**) CCCP-induced autolysis is growth phase-dependent. The graph depicts kinetics of CCCP-induced autolysis for early exponential (OD_600_ =0.2), mid-exponential (OD_600_ =0.6), late-exponential (OD_600_ =1.0) and stationary growth phase (overnight) cells. To rule out a contribution of cell density to the observed autolysis process, the late exponential phase (OD_600_ =1.0) cells were diluted to the same density as early exponential phase cells (OD_600_ 0.2) before CCCP addition. (**d**) CCCP-induced autolysis is neither a programmed cellular response nor induced by prophage activation. The graph depicts the kinetics of CCCP-induced autolysis in *B. subtilis* cells cured of prophages, or in wild-type cells treated with a high concentration (100 µg/ml) of the translation inhibitor chloramphenicol to suppress potential induction of lysis-promoting proteins. See Fig. S3 for additional controls. The graphs depict the mean and SD of OD_600_ measurements performed in technical triplicate. Scale bar, 3 μm. Strains used: *B. subtilis* 168 (wild-type) and *B. subtilis* Δ6 (Δprophage).

The medium and growth phase dependence of CCCP-induced autolysis implies that the process is highly dependent on the cellular metabolic state and likely on corresponding gene expression profiles, and is not an inevitable consequence of PMF collapse. Thus, we were curious whether the observed autolysis represents an example of programmed cell death induced in response to low PMF^37^. To test this, we analysed the lysis of strains lacking individual envelope stress-linked ECF sigma factors (SigM, W, V, X, Y and Z), SigI, which has recently been linked to the regulation of autolysins^20,36–38^, and cells deficient for the membrane stress response proteins PspA and LiaH^20,38–40^. While slight changes in lysis kinetics were observed, none of the analysed mutants suppressed the CCCP-induced autolysis in a substantial manner (Fig. S3). Similar lysis behaviour was also observed for cultures in which cells’ ability to induce stress responses was inhibited by high concentrations of the translation inhibitor chloramphenicol, applied before CCCP treatment (Fig. 2d). Lastly, *B. subtilis* strain cured from prophages^41,42^ also exhibited autolysis in response to CCCP, thus ruling out that activation of dormant phages is responsible for the observed cell lysis (Fig. 2d). In conclusion, lysis induced by PMF collapse does not appear to be a deliberate cellular response but instead represents a failure to adequately downregulate PG hydrolase activity during growth inhibition.

### Actin homologs MreB, Mbl and MreBH are involved in autolysis induced by PMF collapse

The observation that *B. subtilis* stationary growth phase cultures do not undergo autolysis in response to PMF-collapse suggests that active cell growth is a prerequisite. Indeed, PG hydrolase activities are essential for the cell division and cell elongation processes^41,42^, and autolysis induced by cell wall-targeting antibiotics is frequently linked to cell division in *E. coli*^43,44^. To test whether the autolysis induced by CCCP in *B. subtilis* involves active cell division, we inhibited division by three different approaches: (i) depletion of Pbp2B, which allows the cell division machinery to assemble but prevents septum constriction^45^, (ii) deletion of *ftsA*, which significantly lowers the division frequency^46^, and (iii) depletion of FtsZ, which fully prevents the assembly of the cell division machinery^47^. All three approaches lead to cell filamentation caused by inhibition of cell division, but the CCCP-induced cell lysis remained largely unaffected (Fig. 3a-b, Fig. S4a). This argues that neither active cell division nor an assembled cell-division machinery is required for the autolytic process associated with PMF collapse in *B. subtilis*.

**Figure 3:**
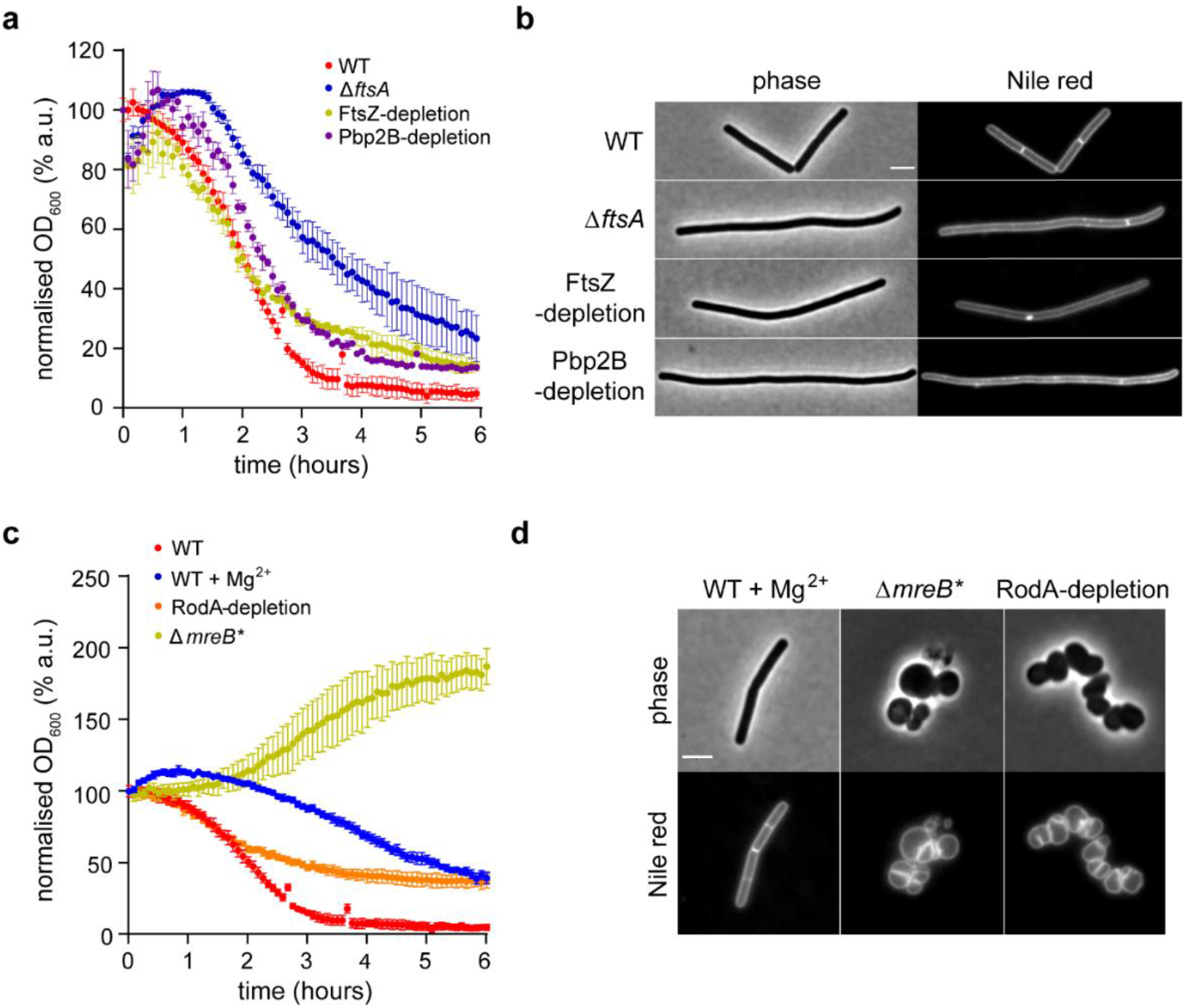
*B. subtilis* autolytic behaviour is linked to the cell elongation process. (**a**) Cells lacking active cell division sites follow CCCP-induced autolysis kinetics that are comparable to wild-type cells. The graph depicts the kinetics of CCCP-induced autolysis in wild-type cells, FtsA-deficient cells that exhibit reduced septation frequency, and cells depleted of the essential division proteins FtsZ or Pbp2B. (**b**) The reduced or eliminated septation for the cell used in panel a was confirmed through fluorescence microscopy. The panel depicts phase contrast and fluorescent images of cells stained with membrane dye Nile red. Note the lack of septa observed in the long filamentous cells depleted for FtsZ or PBP2B, and the low frequency in cells deficient for FtsA. See Fig. S4A for the same cells in the presence of inducers. (**c**) Absence of MreB-homologs (Δ*mreB* Δ*mbl* Δ*mreBH*, labelled as Δ*mreB*\*), but not the absence of rod-shaped morphology, eliminates CCCP-induced autolysis. The graph depicts the kinetics of CCCP-induced autolysis for wild-type cells in the presence and absence of Mg^2+^ supplementation, cells lacking MreB-homologs (Δ*mreB*, Δ*mbl*, Δ*mreBH*) in the presence of Mg^2+^, and cells depleted for the glycosyltransferase RodA. (**d**) The cell morphologies associated with the gene deletions were confirmed using phase contrast and fluorescence microscopy of cells stained with the membrane dye Nile red. The graphs depict the mean and SD of OD_600_ measurements performed in technical triplicate. Scale bar, 3 μm. Strains used: *B. subtilis* 168 (wild type), *B. subtilis* KS109 (*Pspac-ftsZ*), *B. subtilis* KS108 (*Pspac-pbp2B*), *B. subtilis* KS97 (Δ*ftsA*), *B. subtilis* KS60 (Δ*mreB* Δ*mbl* Δ*mreBH*), and *B. subtilis* KS99 (*Pspac-rodA*).

The second major morphogenetic machinery of *B. subtilis* is the elongasome, which is composed of membrane-associated cytoskeletal structures formed by MreB and its homologs Mbl and MreBH, and the associated cell wall synthetic machinery centred around the glycosyltransferase RodA and transpeptidases PbpA and PbpH^16,46^. Spatially guided by antiparallel MreB/Mbl/MreBH polymers^16,48^, the elongasome incorporates new peptidoglycan strands perpendicular to the cell length axis, thereby driving cell elongation^14,47^. While the cytoplasmic MreB and its homologs cannot interact with extracellular PG hydrolase directly, a regulatory link between MreB-homologs and PG hydrolases LytE and CwlO is well established^14,49^. *B. subtilis* strains deficient in MreB-homologs are viable, enabling testing of their role in the autolytic process. However, these mutants lose the rod-shape morphology of wild-type cells^50^, and require high Mg^2+^ concentrations in the medium for growth^51,52^. Consistent with the observation by Tesson *et al*. that Mg^2+^ regulates PG hydrolase activity in *B. subtilis*^17,18,21,51,52^, supplementation of 20 mM MgCl_2_ into the growth medium indeed reduced the level of lysis in response to CCCP (Fig. 3c). While strains deficient for individual MreB homologs exhibited mildly suppressed lysis phenotypes (Fig. S4b), the lack of all three homologs fully abolished lysis induced by CCCP. Instead, a slow increase in OD was observed in the presence of CCCP, corresponding to about one doubling in 6 hours. Due to the altered cell morphology of this strain, it is difficult to determine whether the increase in OD reflects changes in the cells’ optical properties, changes in cell size, or genuine slow growth in the presence of CCCP. However, sustained growth is unlikely as the strain’s minimal inhibitory concentration for CCCP remains unchanged (Table S1).

To test whether the suppression of autolysis is specifically due to the deletion of *mreB*-homologs or due to the lack of elongasome activity and the associated loss of rod-shaped morphology, we inhibited elongasome activity through depletion of the glycosyltransferase RodA, which is essential for the elongasome-associated cell wall synthesis^17,18,53,54^. Despite similar cell morphology (Fig. 3d, Fig. S4a), the strain depleted for RodA retained a strong autolytic response to CCCP (Fig. 3c). Taken together, this argues for a critical and specific role of MreB-homologs in the cellular processes that link PMF collapse to induced autolysis.

### Autolysis is induced by delocalisation of MreB-homologs

We have previously reported that the cellular localisation of MreB, Mbl and MreBH is sensitive to dissipation of membrane potential^55,56^. Therefore, the delocalisation of MreB-homologs upon membrane potential dissipation could be linked to the observed missregulation of PG hydrolases. Mbl has been linked to the regulation of CwlO via FtsEX, while MreB and MreBH facilitate the regulation of LytE via a currently unknown mechanism^14,15,49^. As no suppression of lysis was observed in strains deficient in CwlO or its regulators, FtsEX (Fig. S1), we focused our analysis on MreB and tested whether CCCP-induced delocalisation correlated with the temperature-dependent induction of autolysis.

As expected from its key role in morphogenesis, the localisation of MreB to membrane-associated filaments was identical irrespective of the growth temperature (Fig. 4a and Fig. S5a). However, after incubation with CCCP for 10 mins, a clear temperature-dependent difference in localisation pattern was observed. Under these conditions, MreB exhibited stronger delocalisation to membrane-associated aberrant clusters at 37°C (Fig. S5), which we have previously shown to disturb membrane lipid domain organisation^55^. Upon extended incubation for 30 min, which is likely more relevant to the delayed autolysis process, a loss of MreB membrane association was observed at 37°C. In contrast, the delocalisation of MreB was much more subtle at 30°C (Fig. 4a, c). The CCCP triggered delocalisation of MreB, thus, correlates with the extent of induced autolysis. Consistent with the notion that the cellular role of MreB in regulating autolysis is independent of the elogasome activity, the CCCP-induced delocalisation of MreB was maintained upon depletion of RodA (Fig. S6). Thus, delocalisation of MreB correlates with autolysis, irrespective of whether MreB is engaged with the elongasome complexes that drive active cell wall synthesis.

**Figure 4:**
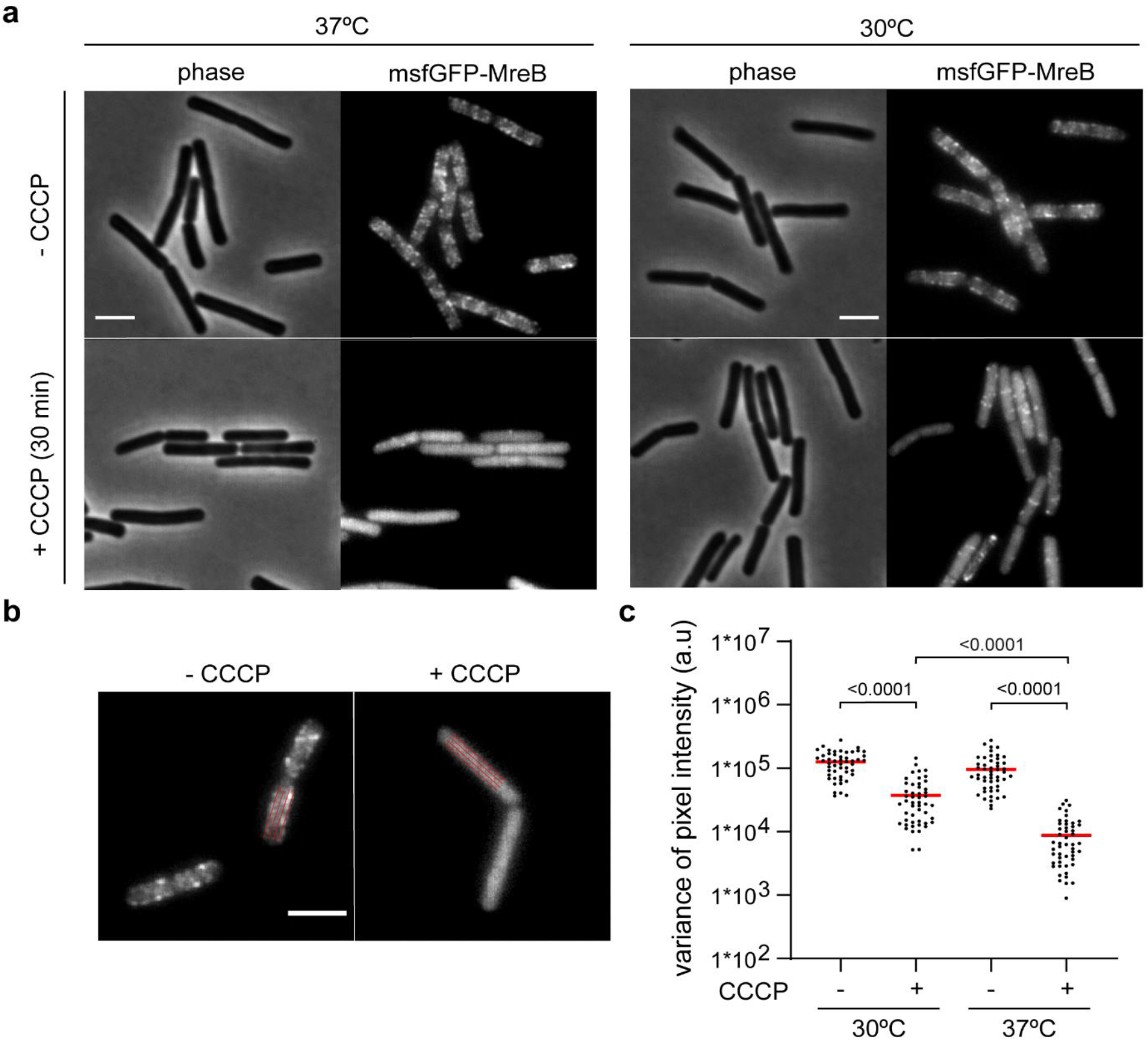
Collapse of PMF triggers delocalisation of MreB in a temperature-dependent manner. (**a**) MreB-polymers dissolve and disassociate from the membrane more extensively upon incubation with CCCP at 37°C. The phase contrast and fluorescence images depict cells grown at 30°C and 37°C, respectively, expressing msfGFP-MreB fusion protein in the presence and absence of CCCP (30 min). Note the CCCP-induced depolymerisation and delocalisation to the cytoplasm that is more extensive at 37°C. See supplementary Fig. S5 for corresponding microscopy images upon 10 min incubation with CCCP. (**b**) A schematic indicates the orientation and thickness of the lines used to measure fluorescence intensity fluctuations, which are used here as a proxy for protein clustering. (**c**) CCCP triggers more extensive depolymerisation of MreB at 37°C. The graph depicts the variance of msfGFP-MreB fluorescence intensity fluctuations for the cells shown in panel a (n=50), together with P-values from an unpaired, two-sided *t*-test. The median variance is indicated with a red line. Scale bar, 3 μm. Strain used: *B. subtilis* HS553 (*msfGFP-mreB*).

### Membrane pore-forming compounds induce lysis through MreB-mediated autolysis in *B. subtilis*

As demonstrated earlier, the protonophore CCCP, which does not directly compromise the membrane permeability barrier, induces autolysis via a cellular pathway involving delocalisation of MreB. However, most membrane-targeting antimicrobial compounds permeabilise the membrane either by forming selective channels or larger, less selective membrane pores^57,58^. This raises the question of whether such compounds induce cell lysis more directly by facilitating leakage of cellular content in a process frequently termed membrananolysis. To test this, we focused on two antimicrobial peptides, each with a different mechanism of membrane permeabilisation: Gramicidin, which forms small cation-specific membrane channels^59^, and the human antimicrobial peptide LL-37, which forms larger, unselective pores^60^. Wild type *B. subtilis* cells treated with gramicidin exhibited slow lysis kinetics, similar to cultures treated with CCCP (Fig. 5a). In contrast, treatment with LL-37 resulted in a much more rapid and extensive collapse of the culture optical density, indicating quick lysis that could be facilitated by extensive leakage of cellular content though large LL-37 pores (Fig. 5a). Surprisingly, both the delayed and rapid lysis induced by gramicidin and LL-37, respectively, were abolished in the strain deficient for major autolysins (Fig. 5b), as well as in the strain that lacks MreB-homologs (Fig. 5c). Crucially, these strains remained growth inhibited (Fig. 5a-c), while the antimicrobial peptides exhibited the same minimal inhibitor concentrations comparable to wild type (Table S1), and retained the ability to fully dissipate the membrane potential (Fig. 5d-f). Together, these findings indicate that, despite their direct membrane-permeabilising activities and the rapid action of LL-37, cell lysis is nonetheless indirect and mediated by PG hydrolases, and is subject to the same MreB-dependent regulatory processes identified for PMF collapse.

**Figure 5:**
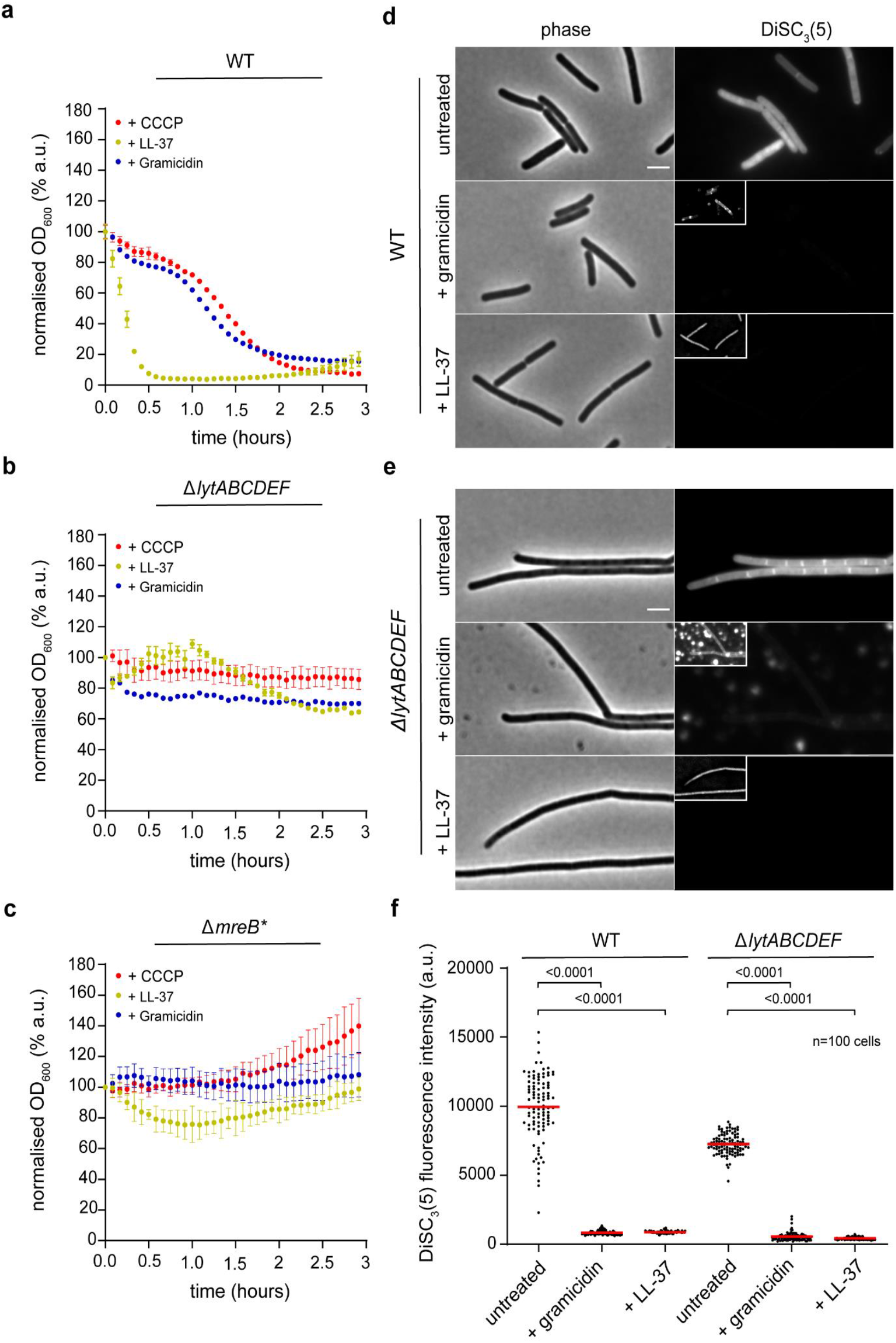
MreB-dependent autolysis is responsible for the bacteriolytic activity of membrane-targeting antimicrobials in *B. subtilis*. Small cation channel gramicidin induces delayed autolysis in a manner comparable to the proton carrier CCCP, while the large pore-forming antimicrobial peptide LL-37 induces more rapid lysis. The graphs depict lysis kinetics in the presence of CCCP, gramicidin and LL-37 for (**a**) the wild-type cells, (**b**) for cells deficient for several autolytic enzymes, and (**c**) for cells deficient for MreB-homologs (Δ*mreB* Δ*mbl* Δ*mreBH*, labelled as Δ*mreB*\*). The graphs depict the mean and SD of OD600 measurements performed in technical triplicate. (**d-e**) Both gramicidin and LL-37 retain their ability to trigger membrane depolarisation in the absence of autolysis. The phase contrast and fluorescence images depict (d) wild type and (e) Δ*lytABCDEF* cells stained with the membrane potential sensitive dye DiSC_3_(5) in the presence and absence of gramicidin and LL-37, respectively. (**f**) Quantification of gramicidin and LL-37-induced membrane depolarisation. The graph depicts DiSC_3_(5) fluorescence levels for the cells (n=100) shown in panels d and e. The median intensity values are indicated with a red line, while the P values represent those of an unpaired, two-sided t-test. Scale bar, 3 μm. Strains used: *B. subtilis* 168 (wild type), *B. subtilis* KS19 (Δ*lytABCDEF*) and *B. subtilis* KS60 (Δ*mreB* Δ*mbl* Δ*mreBH*).

## DISCUSSION

Due to their potential to degrade the peptidoglycan cell wall, careful regulation of autolytic enzyme activity is critical for bacterial cells. However, our understanding of the regulatory processes governing autolytic activity is limited. Arguably, the best-characterised autolytic enzyme regulatory systems are those involving teichoic acid-dependent regulation in *Streptococcus pneumoniae*^22,61^, and those involving the FtsEX-complex^6,14,62–64^. The behaviour of *B. subtilis* in response to energy starvation and low PMF is one of the earliest characterised examples of bacterial autolysis^25–27^, yet the mechanism that underpins the process has remained enigmatic. To date, the prevailing model put forward by Jolliffe *et al* argues that PMF and pH-dependent membrane surface properties inhibit autolytic enzyme activity in its proximity^26^. Here, we postulate that, rather than relying on biochemical processes linked to pH, the autolysis induced by PMF collapse is a more complex cellular process involving the MreB cytoskeleton and its delocalisation upon membrane potential dissipation. Due to the many pleiotropic consequences of PMF collapse and, beyond FtsEX, our relatively superficial understanding of autolysin regulators in *B. subtilis*, it is difficult to speculate about the molecular mechanisms linking the intracellular localisation of the MreB cytoskeleton to extracellular control of autolytic enzyme activity. However, we have previously shown that the small cyclic antimicrobial peptide cWFW induces delocalisation of MreB and associated autolysis under conditions where cWFW only induces mild membrane depolarisation^11^. Moreover, expression of the native and non-membrane depolarising toxin BsrG also triggers delocalisation of MreB and autolysis^63^. In this case, deleting either *mreB* or *lytD* is fully sufficient to abolish autolysis. Together, these findings suggest that delocalisation of MreB, rather than a collapse of PMF, is the critical autolysis-initiating step (Fig. 6). Moreover, the observation that this property is not linked to MreB’s role in cell wall synthesis is intriguing as it argues for the dual, functionally uncloupled role of the MreB-cytoskeleton in regulating peptidoglycan biosynthesis and its turnover.

**Figure 6:**
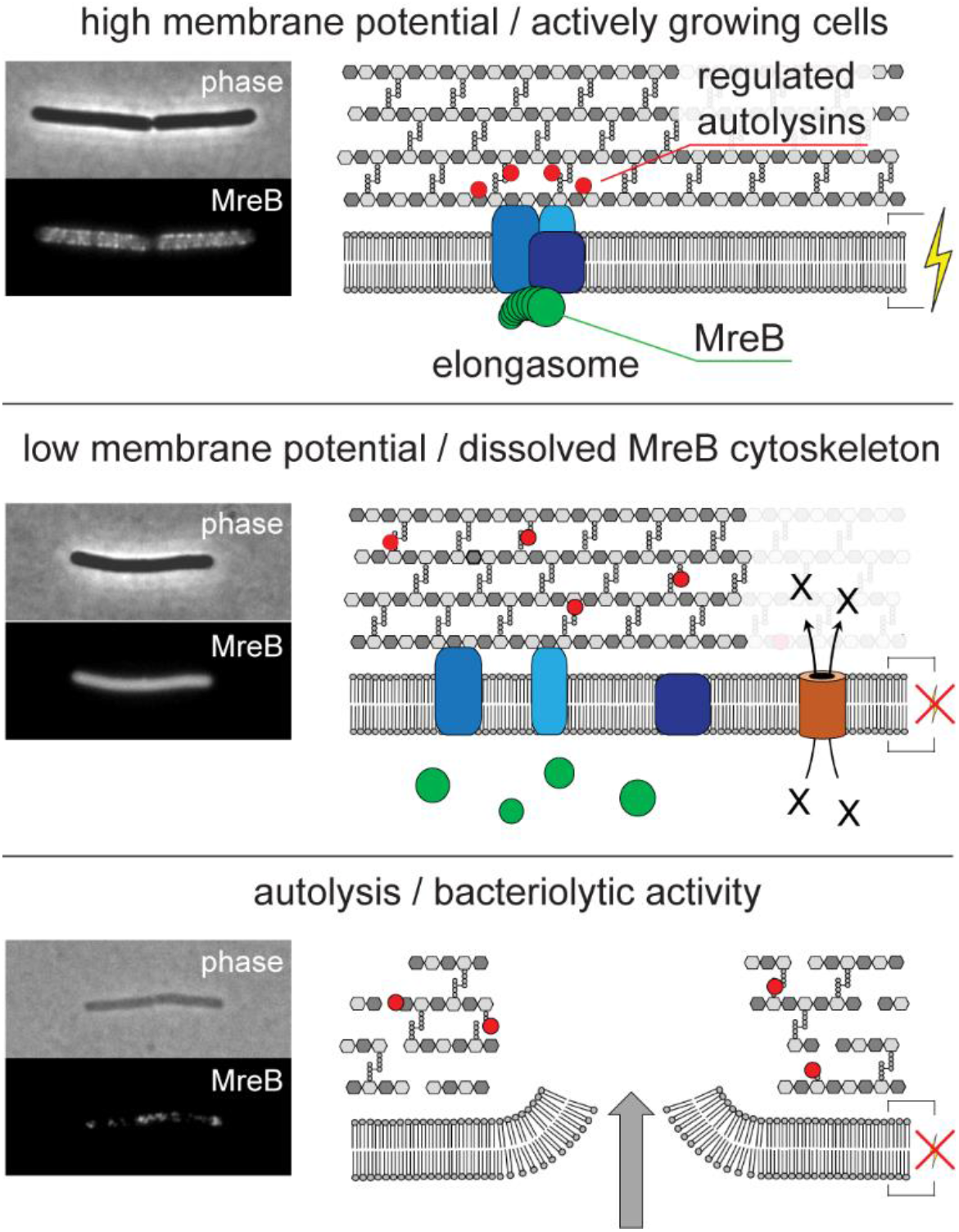
Schematic model of MreB delocalisation–induced autolysis. In well-energised cells, MreB filaments engage with the cell wall synthesis machinery and guide insertion of new peptidoglycan strands into an existing cell wall matrix. Under these conditions, autolysins are tightly controlled through an unknown mechanism but in a manner that depends on the membrane association of MreB filaments (upper panel). Upon extended dissipation of the membrane potential, MreB dissociates from the membrane, leading to dysregulation of autolysins and runaway degradation of the cell wall (middle panel). Ultimately, the degradation of the cell wall leads to areas that are unable to support the membrane under turgor, resulting in leakage of cytoplasmic contents and cell disintegration (lower panel).

The finding that the rapid lysis induced by the large pore-forming peptide LL-37 is also autolytic in nature is striking. Membrane-targeting antibiotics and antimicrobial peptide studies typically centre on the compound’s ability to induce membrane permeabilisation. This property is relevant for understanding the molecular basis of antibacterial activity, but also for evaluating the potential cytotoxic activity that is frequently linked to membrane permeabilisation^65^. Indeed, whether membrane pore formations occur for the membrane-targeting last resort antibiotic daptomycin has been a matter of considerable debate, largely driven by disagreeing *in vivo* and *in vitro* results^66^. Recently, we reported that the apparent pore formation observed with daptomycin in *B. subtilis* is caused by autolysis^67^. Daptomycin thus triggers membrane permeabilisation in a manner that is indirect and not reflecting its intrinsic activity towards bacterial membranes. Our findings with the human antimicrobial peptide LL-37 presented here, as well as for the membrane-acting antiseptic octenidine published elsewhere^68^, highlight the need to consider autolytic processes even when the compound’s direct antibacterial mechanism of action is membrane permeabilisation. Moreover, they demonstrate that the ability to induce extensive membrane permeabilisation and substantial leakage of cytoplasmic content does not necessarily translate to a decline in culture optical density, or loss of phase contrast on microscopic observation. Hence, equating molecular processes such as membrane pore formation with cellular phenomena such as cell lysis is not as trivial as frequently assumed.

## METHODS

### Strains and growth conditions

All strains used in this study are listed in Supplementary Table 2, and were routinely maintained on nutrient agar (NA) (Oxoid) with supplements. Selective media contained chloramphenicol (5 μg/ml), erythromycin (1 μg/ml), kanamycin (2 and 5 μg/ml), tetracycline (6 μg/ml), spectinomycin (50 μg/ml), phleomycin (1 μg/ml), ampicillin (100 μg/ml), IPTG (0.1-1 mM) and xylose (0.1-0.5 %). Membrane-active compounds were used at concentrations of gramicidin (10 μg/ml), LL-37 (40 μg/ml), CCCP (carbonyl cyamide 3-chlorophenylhydrazone; 100 μM). Mg_2_SO_4_ (20 mM) supplement was required for all overnight cultures and experiments with strains deleted for *mre* homologs.

### DNA manipulation and strain constructions

All *B. subtilis* transformations were carried out via natural competence as previously described^69^. DNA for the transformation was provided either as genomic DNA purified from the respective *B. subtilis* strain or as linearised plasmid DNA containing the desired genetic construct. Details of individual strain constructions are listed in Supplementary Table 2.

### Lysis kinetic assay

To measure lysis kinetics, cells were grown overnight in LB at 30°C without antibiotics, diluted into fresh medium, and allowed to reach mid-exponential growth phase at 30°C or 37°C unless specified otherwise. Strains were transferred into a transparent 96-well microplate (Greiner Bio-One) in technical triplicate before the addition of the membrane-active compounds. If needed, the cells were incubated with the ribosome inhibitor chloramphenicol (100 μg/ml) to inhibit protein translation for 30 min before adding the membrane-active compounds. Samples were assayed in a FLUOstar OPTIMA or SPECTROstar Nano (BMG Labtech) with orbital agitation at 200 rpm, and OD_600_ was measured every 5 min. Data obtained were then normalised against the initial OD upon addition of test compounds. For depletion experiments, strains were assayed as stated above, but with the following modifications. Cells were grown overnight in LB supplemented with inducer, followed by resuspension in fresh medium containing inducer the following morning. To ensure robust growth, cells were allowed to undergo three doublings before being washed and resuspended in inducer-free medium. The growth and lysis experiments in media with altered carbon sources were carried out in SMM-based media with composition as detailed elsewhere^70^.

### Cell viability assay

For cell viability measurement, the cells were grown and treated as described for the lysis kinetic assay at 37°C. Samples were taken hourly and diluted 1000-fold in LB medium to dilute the membrane-active compound below inhibitory levels. 1:10 dilution series were prepared, spotted onto nutrient agar plates, and incubated overnight at 30°C. For control, untreated samples were collected before the addition of test compounds and prepared as described above.

### Minimum inhibitory concentration assay

For the determination of MIC, cells were grown overnight in LB at 30°C before resuspension in fresh medium the following day and allowed to undergo at least three doublings to the mid-exponential growth phase at 37°C. A 1:2 serial dilution of the membrane-active compounds was prepared in a clear 96-well microplate (Greiner Bio-One) in technical triplicate, followed by the addition of cells to a final inoculum of 10000 cells/ml. The microtiter plates were incubated for 16 hours with shaking at 37°C. The highest concentration of the test compounds at which no visually detectable growth was observed was determined as the MIC.

### General fluorescent microscopy

For the fluorescent microscopy, the cells were grown and treated as described for the lysis kinetic assay at 30°C or 37°C. When indicated, cell suspensions were stained for 15 min with 100 nM SYTOX Green, 2 μM DiSC_3_(5) (3,3’-Dipropylthiadicarbocyanine Iodine), 0.5 μg/ml Nile Red or 2 μg/ml FM 5-95 (N-(3-Trimethylammoniumpropyl)-4-(6-(4(Diethylamino)phenyl)hexatrienyl) Pyridinium Dibromide) all from ThermoFisher Scientific. The cells were immobilised onto 1.2% agarose in H_2_O spread thinly onto microscope slides as previously described^51^. Microscopy was carried out with Nikon Eclipse Ti equipped with Nikon Plan Apo 100x/1.40 Oil Ph3 objective, Photometrics Prime sCMOS camera, and either Lambda LS (Sutter Instrument) or CoolLed pE-300 (CoolLED) light source. Metamorph 7.7 were used to acquire images.

### Image analysis

Cellular SYTOX green and DiSC_3_ fluorescence signals were quantified from background-subtracted micrographs using Fiji^71^.

### Statistical analyses

All experiments were carried out in at least biological duplicates. The averages and standard deviations, as well as the statistical significance calculations, were performed using GraphPad Prism.

## Author contributions

KHS and AK performed the experiments, KHS, AK and HS analysed data, AK and HS wrote the paper, and HS acquired the funding for the research.

## Data Availability

Source data for all figures and graphs presented in the manuscript will be made available via Newcastle University’s research data repository once the manuscript takes its final, peer-reviewed form. The strains and plasmids are available upon request to H.S.

## Acknowledgements

We acknowledge James Donahue for constructing strains and Richard Daniel for valuable comments during the manuscript preparation.

