## Supplementary Information for "Membrane-targeting antimicrobials trigger lysis in *Bacillus subtilis* by disturbing the MreB-dependent regulation of peptidoglycan hydrolases"

Authors: Kenneth H. Seistrup[1, 2], Alan Koh[1,3,4], and Henrik Strahl[1]

Affiliation:

1: Centre for Bacterial Cell Biology, Biosciences Institute, Faculty of Medical Sciences, Newcastle University, Newcastle upon Tyne, UK

Current affiliations:

2: Molecular Devices, San José, US

3: MRC Laboratory of Medical Sciences, London, UK

4: Institute of Clinical Sciences, Imperial College, London, UK

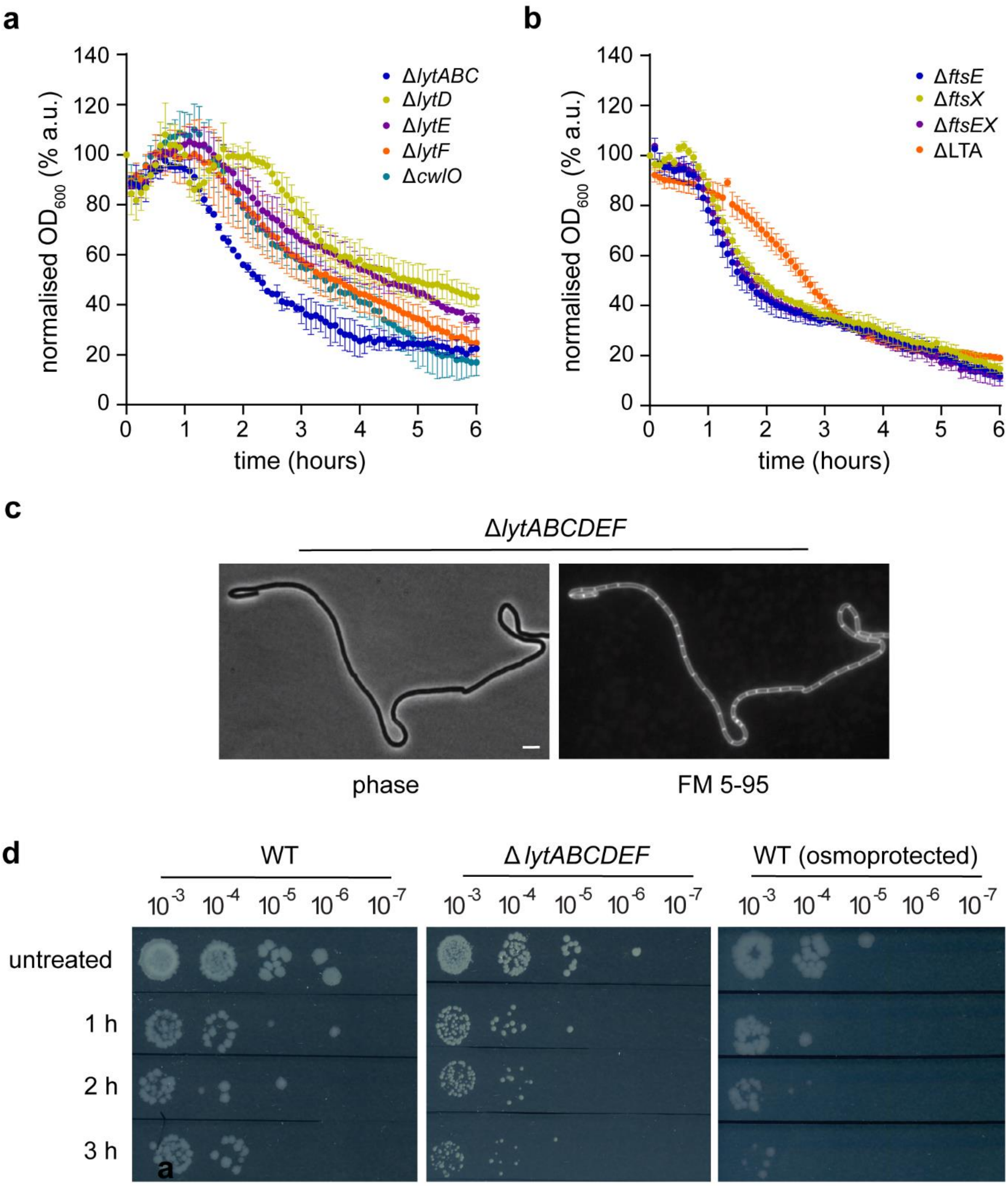


**Figure S1: CCCP-induced autolysis process involves several PG hydrolases.**

(**a**) None of the *B. subtilis* main PG hydrolases is solely responsible for the CCCP-induced autolysis. The graph shows the kinetics of CCCP-induced autolysis for *B. subtilis* strains lacking the PG hydrolases LytC, LyD, LytE, LytF or CwlO. (**b**) Neither the FtsEX complex nor lipoteichoic acids are involved in the CCCP-induced autolysis. The graph shows the kinetics of CCCP-induced autolysis in B. subtilis strains lacking FtsE, FtsX, or both, and deficient in lipoteichoic acid synthesis. The data points in both graphs represent the mean and SD of $OD_{600}$ measurements performed in technical triplicate. (**c**) *B. subtilis* strain deleted for *lytACB, lytD, lytE* and *lytF* exhibits a strong morphology defect caused by suppressed cell separation, resulting in growth as long chains of cells. The images depict phase contrast and fluorescence microscopy of cells stained with the membrane dye FM 5-95. Note the presence of completed septa, indicating that cell division is not inhibited in these strains. Scale bar, 3 µm. (**d**) CCCP acts bactericidally despite the lack of lysis. A comparable reduction in cell viability was observed after treatment with CCCP, even when lysis was suppressed by deletion of *lytABCDEF* or by incubation in an osmoprotective medium (LB + 0.5 M sucrose). Strains used: *B. subtilis* 168 (wild type), *B. subtilis* KS10 (Δ*lytABC*), *B. subtilis* KS9 (Δ*lytD*), *B. subtilis* KS8 (Δ*lytE*), *B. subtilis* KS7 (Δ*lytF*), *B. subtilis* KS6 (Δ*cwlO*), *B. subtilis* KS104 (Δ*ftsX*), *B. subtilis* KS105 (Δ*ftsE*), *B. subtilis* KS107 (Δ*ftsEX*), *B. subtilis* AK066B (ΔLTA) and *B. subtilis* Δ6 (Δ*prophages*).

**a**

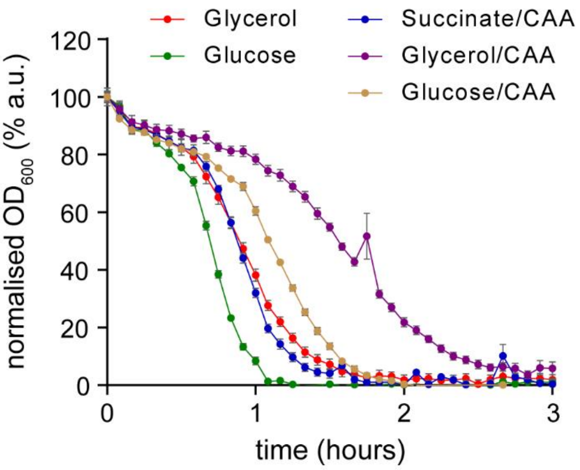


**b**

| carbon source | growth rate (1/h) | lysis rate (1/h) |
|---|---|---|
| Glycerol | 1.1 | 2.8 |
| Glucose | 1.2 | 5.8 |
| Succinate + CAA | 1.3 | 3.6 |
| Glycerol + CAA | 1.4 | 1.6 |
| Glucose + CAA | 1.9 | 2.8 |

**c**

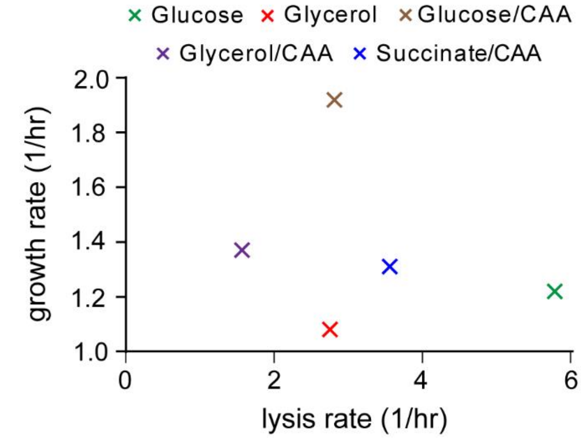


**Figure S2: Rate of CCCP-induced autolysis does not correlate with growth rate.**

(**a**) Rate of lysis depends on the carbon source and casaminoacids (CAA) supplementation in an SMM-based medium. The graph shows the kinetics of CCCP-induced autolysis in *B. subtilis* grown in different media, represented as the mean and SD of $OD_{600}$ measurements performed in technical triplicate. (**b**) The medium carbon source and CCCP supplementation modulate both growth and lysis rates. The values were calculated for logarithmically growing cells prior to CCCP addition, and the maximal lysis rates observed after CCCP addition. No correlation was observed between growth and lysis rates. Strain used: *B. subtilis* 168 (wild type).

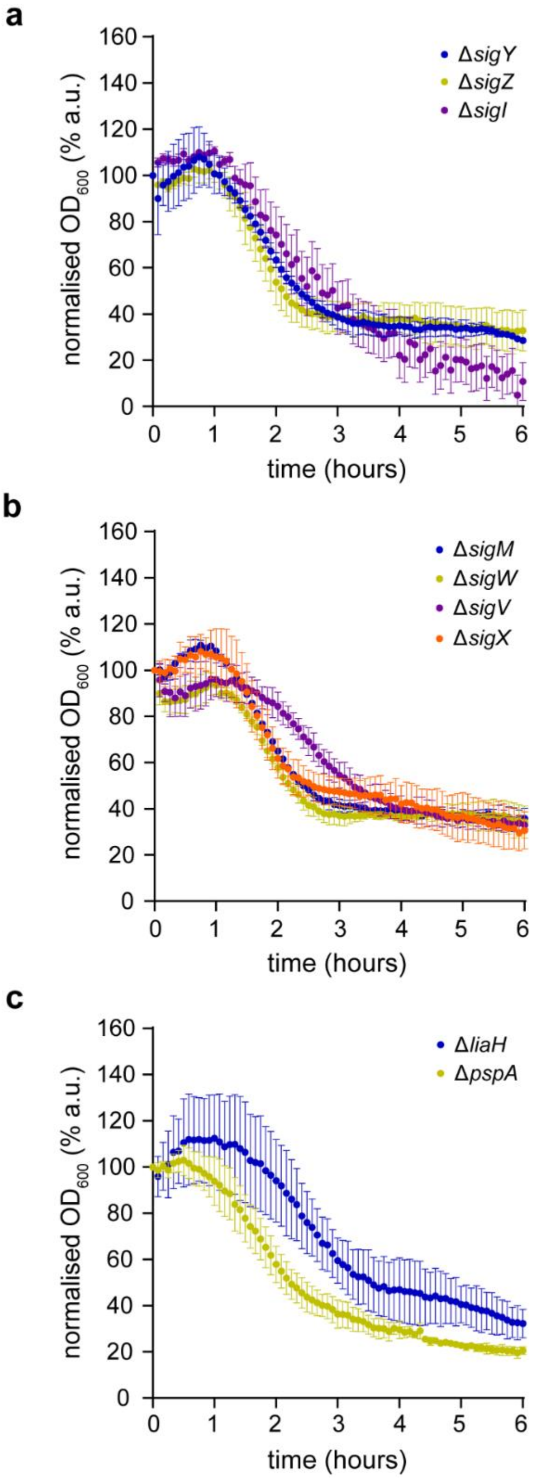


**Figure S3: CCCP-induced autolysis is not a programmed cellular response.**

(a-b) Lack of individual extracytoplasmic function (ECF) sigma-factors of SigI does not significantly suppress the CCC-induced autolytic process. (b) Induction of the envelope stress response proteins LiaH and PspA is also not implicated in the CCC-induced autolysis. Strains used: B. subtilis 168 (wild type), B. subtilis KS44 (Δ*sigY*), B. subtilis KS41 (Δ*sigM*), B. subtilis KS43 (Δ*sigW*), B. subtilis KS49 (Δ*sigZ*), B. subtilis KS48 (Δ*sigV*), B. subtilis KS42 (Δ*sigX*), B. subtilis KS121 (Δ*sigI*), B. subtilis KS4 (Δ*pspA*) and B. subtilis KS5 (Δ*liaH*).

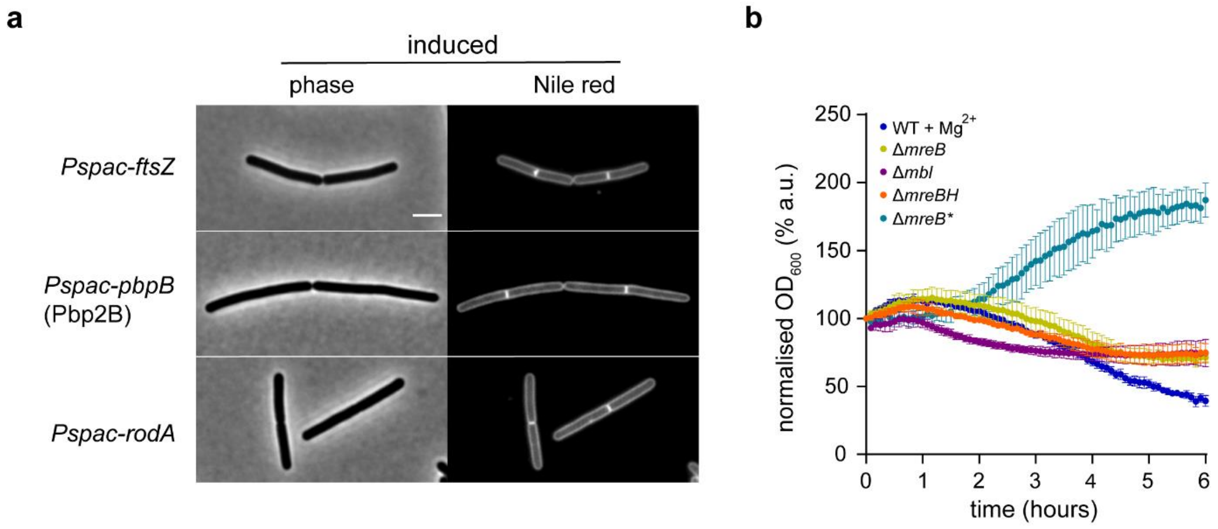


**Figure S4: Morphogenetic processes and genes involved in CCCP-induced autolysis.**

(a) Morphology of strains encoding inducible copies of ftsZ, pbpB and rodA in the presence of the inducer IPTG. The panel depicts phase contrast and fluorescent images of cells stained with the membrane dye Nile red. See Figure 3 b for the same cells in the absence of inducers. (b) Absence of all three MreB-homologs (Δ*mreB* Δ*mbl* Δ*mreBH*, labelled as Δ*mreB**) is required for suppressing CCCP-induced autolysis. The graph depicts the kinetics of CCCP-induced autolysis for wild-type cells in the presence and absence of $Mg^{2+}$ supplementation, and cells lacking individual MreB-homologs in the presence of $Mg^{2+}$. Strains used: *B. subtilis* 168 (wild type), *B. subtilis* KS109 (*Pspac-ftsZ*), *B. subtilis* KS108 (*Pspac-pbpB*), *B. subtilis* KS99 (*Pspac-rodA*), *B. subtilis* KS36 (Δ*mreB*), *B. subtilis* KS37 (Δ*mbl*) and *B. subtilis* KS38 (Δ*mreBH*).

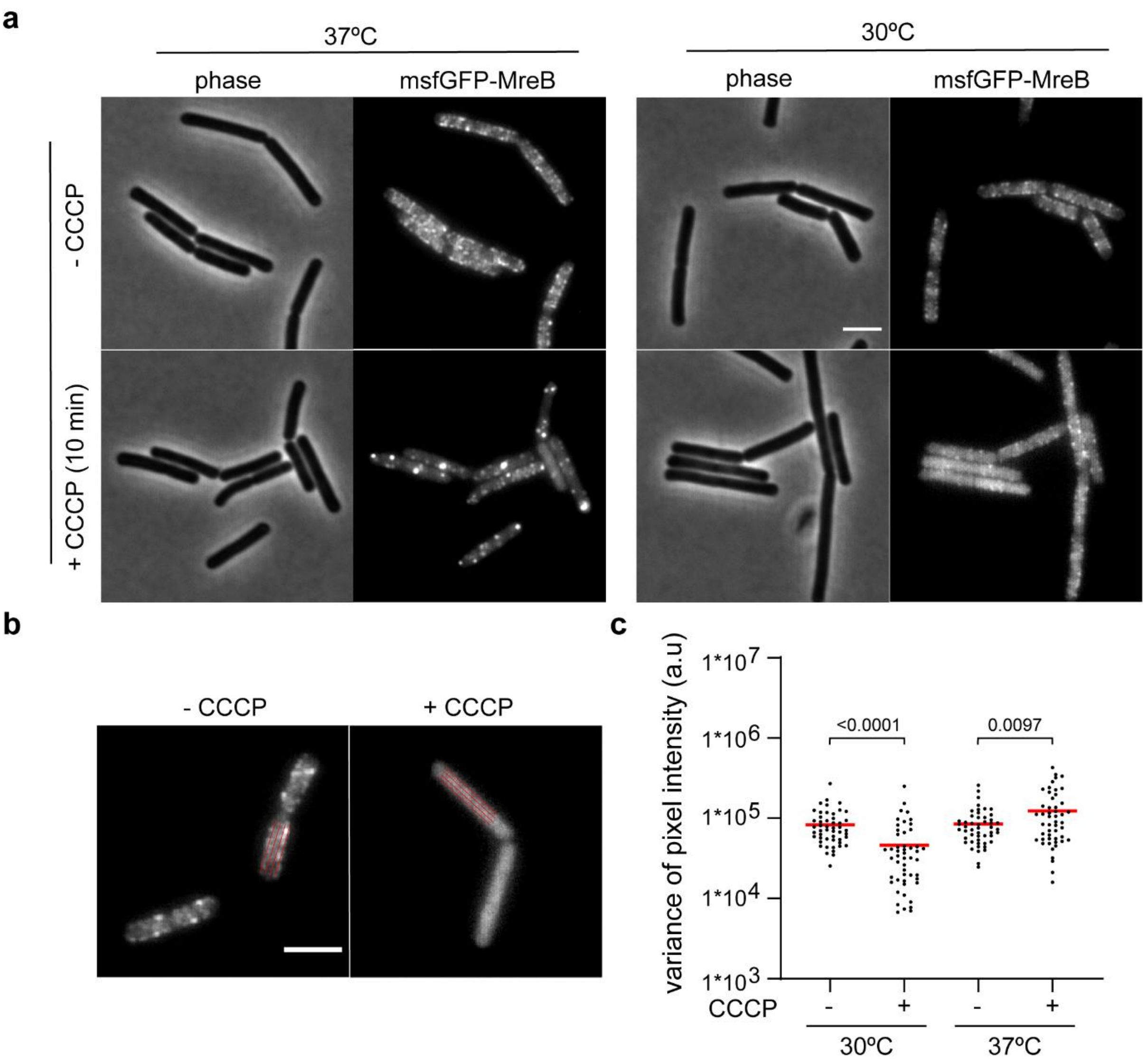


**Figure S5: Clustering at the membrane precedes the membrane dissociation of MreB induced by CCCP at 37°C.**

(a) Unlike at later stages associated with diffuse and cytoplasmic localisation of MreB (Figure 4), shorter (10 min) incubation with CCCP induces clustering of MreB at the membrane. The phase contrast and fluorescence images depict cells grown at 30°C and 37°C, respectively, and expressing the msfGFP-MreB -fusion protein in the presence and absence of CCCP (10 min). (**b**) A schematic indicates the orientation and thickness of the lines used to measure fluorescence intensity fluctuations, which are used here as a proxy for protein clustering. (**c**) MreB remains clustered, indicating the presence of MreB-polymers, upon 10 min incubation with CCCP, irrespective of the growth temperature. The graph depicts the variance of msfGFP-MreB fluorescence intensity fluctuations for the cells shown in panel a (n=50), together with P-values from an unpaired, two-sided *t*-test. The median variance is indicated with a red line. Scale bar, 3 µm. Strain used: *B. subtilis* HS553 (*msfGFP-mreB*).

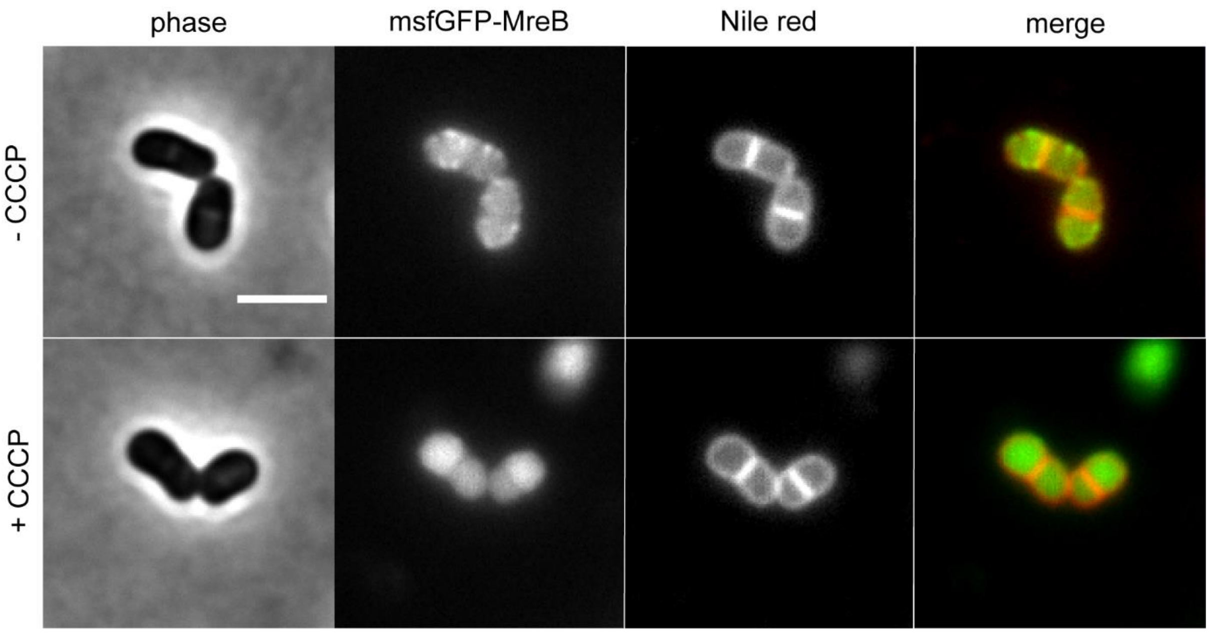


**Figure S6: MreB retains CCCP-induced delocalisation behaviour in the absence of elongasome activity.**

The panel depicts phase-contrast and fluorescent images of cells encoding msfGFP-MreB upon RodA depletion and staining with the membrane dye Nile red. Note the CCCP-induced (10 min incubation at 37°C) depolymerisation and dissociation of MreB from the membrane under conditions in which elongasome activity is inhibited through depletion or RodA, thus disconnecting MreB from active cell wall synthesis. Strain used: B. subtilis KS101 (*Pspac-rodA, msfGFP-mreB*). Scale bar, 3 µm

**Table S1: Minimum inhibitory concentrations of used compounds**

| strain | relevant genotype | medium | CCCP (µM) | Gramicidin (µM) | LL-37 (µM) |
|---|---|---|---|---|---|
| *B. subtilis* 168 | wild type | LB | 12 | 0.6 | 2.25 |
| *B. subtilis* 168 | wild type | LB + 500 mM sucrose | 12 | 2.5 | 4.5 |
| *B. subtilis* 168 | wild type | LB + 20 mM $MgSO_4$ | 6 | nd | nd |
| *B. subtilis* 168 | Δ*lytABCDEF* | LB | 12 | 0.3 | 2.25 |
| *B. subtilis* KS60 | Δ*mreB* Δ*mbl* Δ*mreBH* | LB + 20 mM MgSO4 | 6 | nd | nd |

nd, not determined

**Table S2: Strains used in this study**

| strain | relevant genotype[a] | construction[b] | induction | source |
|---|---|---|---|---|
| *B. subtilis* 168 | *trpC2* (wild type) | - | - | [1] |
| *B. subtilis* 1A792 | *lytABC::kan lytD::tet lytE::cat lytF::spc* | - | - | [2] |
| *B. subtilis* KS19 | *lytABC::kan lytD::tet lytE::cat lytF::spc* | 1A792→168 | - | this work |
| *B. subtilis* KS10 | *lytABC::kan* | 1A792→168 | - | this work |
| *B. subtilis* KS9 | *lytD::tet* | 1A792→168 | - | this work |
| *B. subtilis* KS8 | *lytE::cat* | 1A792→168 | - | this work |
| *B. subtilis* KS7 | *lytF::spc* | 1A792→168 | - | this work |
| *B. subtilis* PDC463 | *cwlO::spc* | - |  | [3] |
| *B. subtilis* KS6 | *cwlO::spc* | BP079→168 | - | this work |
| *B. subtilis* 4501 | *ftsX::kan* | - | - | [3] |
| *B. subtilis* KS104 | *ftsX::kan* | 4501→168 | - | this work |
| *B. subtilis* 4503 | *ftsE::kan* | - |  | [3] |
| *B. subtilis* KS105 | *ftsE::kan* | 4503→168 | - | this work |
| *B. subtilis* PDC484 | *ftsEX::spc* | - |  | [3] |
| *B. subtilis* KS107 | *ftsEX::spc* | PDC484→168 | - | this work |
| *B. subtilis* AG600 | *ltaS::cat yfnI::erm yqgS::spc* | - | - | [4] |
| *B. subtilis* AK066B | *ltaS::cat yfnI::erm yqgS::spc* | AG600K→168 | - | this work |
| *B. subtilis* HB10216 | *sigM::kan* | - | - | [5] |
| *B. subtilis* KS41 | *sigM::kan* | HB10216→168 | - | this work |
| *B. subtilis* HB7007 | *sigX::spc* | - | - | [5] |
| *B. subtilis* KS42 | *sigX::spc* | HB7007→168 | - | this work |
| *B. subtilis* HB4246 | *sigW::erm* | - | - | [6] |
| *B. subtilis* KS43 | *sigW::erm* | 1A905→168 | - | this work |
| *B. subtilis* HB4245 | *sigY::erm* | - | - | [6] |
| *B. subtilis* KS44 | *sigY::erm* | 1A909→168 | - | this work |
| *B. subtilis* HB0028 | *sigV::kan* | - | - | [7] |
| *B. subtilis* KS48 | *sigV::kan* | HB0028→168 | - | this work |
| *B. subtilis* HB0032 | *sigZ::kan* | - | - | [7] |
| *B. subtilis* KS49 | *sigZ::kan* | HB0032→168 | - | this work |
| *B. subtilis* 4265 | *sigI-rsgI::kan* | - | - | [8] |
| *B. subtilis* KS121 | *sigI-rsgI::kan* | 4265→168 | - | this work |
| *B. subtilis* BKE06180 | *pspA::erm* | - | - | [9] |
| *B. subtilis* KS4 | *pspA::erm* | BKE06180→168 | - | this work |
| *B. subtilis* BKE33120 | *liaH::erm* | - | - | [9] |

| | | | | |
|---|---|---|---|---|
| *B. subtilis* KS5 | *liaH::erm* | BKE33120→168 | - | this work |
| *B. subtilis* 3728 | *Ωneo3427 ΔmreB* | - | - | [10] |
| *B. subtilis* KS36 | *Ωneo3427 ΔmreB* | 3728→168 | - | this work |
| *B. subtilis* 4261 | *mbl::cat* | - | - | [8] |
| *B. subtilis* KS37 | *mbl::cat* | 4261→168 | - | this work |
| *B. subtilis* 4262 | *mreBH::erm* | - | - | [8] |
| *B. subtilis* KS38 | *mreBH::erm* | 4262→168 | - | this work |
| *B. subtilis* KS69 | *amyE::spc Pxyl-msfGFP-mreB* | - | 0.5% xylose | [11] |
| *B. subtilis* YK2245 | *rodA::Pspac-rodA* (*kan*) | - | 0.1-1 mM IPTG | [12] |
| *B. subtilis* KS101 | *rodA::Pspac-rodA (kan)*<br>*amyE::spc Pxyl-msfGFP-mreB* | pJS105→KS99 | 0.5% xylose<br>0.1-1 mM IPTG | this work |
| *B. subtilis* Δ6 | ΔSPβ sublancin 168-sensitive Δ*skin* ΔPBSX Δprophage1 *pks::cat* Δprophage 3 (*cat*) | - | - | [13] |
| *B. subtilis* 1801 | *ftsZ::Pspac-ftsZ* (*phl*) | - | 0.1-1 mM IPTG | [14] |
| *B. subtilis* KS109 | *ftsZ::Pspac-ftsZ* (*phl*) | 1801→168 | 0.1-1 mM IPTG | this work |
| *B. subtilis* 799 | *ftsL::Pspac-pbp2B* (*kan*) | - | 0.1-1 mM IPTG | [15] |
| *B. subtilis* KS108 | *trpC2 ftsL::Pspac-pbp2B* (*kan*) | 799→168 | 0.1-1 mM IPTG | this work |
| *B. subtilis* YK206 | *purA16 metB5 hisA3 guaB ftsA::erm* | - | - | [16] |
| *B. subtilis* KS97 | *ftsA::erm* | YK206→168 | - | this work |
| *B. subtilis* 4277 | *Ωneo3427 ΔmreB mbl::cat mreBH::erm rsgI::spc* | - | - | [8] |
| *B. subtilis* KS60 | *Ωneo3427 ΔmreB mbl::cat mreBH::erm rsgI::spc* | 4277→168 | - | [17] |
| *B. subtilis* JS06 | *ponA::cat* | - | - | [18] |
| *B. subtilis* KS95 | *ponA::cat* | JS06→168 | - | this work |
| *B. subtilis* HS553 | *mreB::msfGFP-mreB* | - | - | this work |

[a] *kan*, kanamycin resistance; *ery*, erythromycin resistance; *cat*, chloramphenicol resistance; *spc*, spectinomycin resistance; *tet*, tetracycline resistance; *phl*, phleomycin resistance.

[b] genomic DNA of the indicated strain transformed into the recipient strain
